# Comprehensive cross-disorder analyses of *CNTNAP2* suggest it is unlikely to be a primary risk gene for psychiatric disorders

**DOI:** 10.1101/363846

**Authors:** Claudio Toma, Kerrie D. Pierce, Alex D. Shaw, Anna Heath, Philip B. Mitchell, Peter R. Schofield, Janice M. Fullerton

## Abstract

The contactin-associated protein-like 2 *(CNTNAP2)* gene is a member of the neurexin superfamily. *CNTNAP2* was implicated in the cortical dysplasia-focal epilepsy (CDFE) syndrome, a recessive disease characterized by intellectual disability, epilepsy, language impairments and autistic features. Associated SNPs and heterozygous deletions in *CNTNAP2* have also frequently been reported in autism, schizophrenia and other psychiatric or neurological disorders. We aim here to gain conclusive evidence for the role of *CNTNAP2* in susceptibility to psychiatric disorders by the comprehensive analysis of large genomic datasets. In this study we used: i) summary statistics from the Psychiatric Genomics Consortium (PGC) GWAS; ii) examined all reported *CNTNAP2* structural variants in patients and controls; iii) performed cross-disorder analysis of functional or previously associated SNPs; iv) and conducted burden tests for pathogenic rare variants using sequencing data (4,483 ASD and 6,135 schizophrenia cases, and 13,042 controls).

In a CNV mircroarray study, we previously identified a 131kb deletion in *CNTNAP2* intron 1, removing a FOXP2 transcription factor binding site in an extended BD family. Here we perform a quantitative-PCR validation showing imperfect segregation with disease (5 bipolar disorder relatives). The distribution of CNVs across *CNTNAP2* in psychiatric cases from previous reports was no different from controls of the database of genomic variants. Gene-based association testing did not implicate common variants in autism, schizophrenia or other psychiatric phenotypes. The association of proposed functional SNPs rs7794745 and rs2710102, reported to influence brain connectivity, was not replicated; nor did functional SNPs yield significant results in meta-analysis across psychiatric disorders. Disrupting *CNTNAP2* rare variant burden was not higher in autism or schizophrenia compared to controls. This large comprehensive candidate gene study indicates that *CNTNAP2* may not be a robust risk gene for psychiatric phenotypes.

**AUTHOR SUMMARY:** Genetic mutations that disrupt both copies of the *CNTNAP2* gene lead to severe disease, characterized by profound intellectual disability, epilepsy, language difficulties and autistic traits. Researchers hypothesized that this gene may also be involved in autism given some overlapping clinical features with this disease. Indeed, several large DNA deletions affecting one of the two copies of *CNTNAP2* were found in some patients with autism, and later also in patients with schizophrenia, bipolar disorder, ADHD and epilepsy, suggesting that this gene was involved in several psychiatric or neurologic diseases. Other studies considered genetic sequence variations that are common in the general population, and suggested that two such sequence variations in *CNTNAP2* predispose to psychiatric diseases by influencing the functionality and connectivity of the brain. In the current study, we report the deletion of one copy of *CNTNAP2* in a patient with bipolar disorder from an extended family where five relatives were affected with this condition. To better understand the involvement of *CNTNAP2* in risk of mental illness, we performed several genetic analyses using a series of large publically available or in-house datasets, comprising many thousands of patients and controls. Despite the previous consideration of *CNTNAP2* as a strong candidate gene for autism or schizophrenia, we show that neither common, deletion nor ultra-rare variants in *CNTNAP2* are likely to play a major role in risk of psychiatric diseases.

## INTRODUCTION

The contactin-associated protein-like 2 (*CNTNAP2*) is located on chromosome 7q35-36.1, and consists of 24 exons spanning 2.3Mb, making it one of the largest protein coding genes in the human genome. This gene encodes the CASPR2 protein, related to the neurexin superfamily, which localises with potassium channels at the juxtaparanodal regions of the Ravier nodes in myelinated axons, playing a crucial role in the clustering of potassium channels required for conduction of axon potentials [1]. *CNTNAP2* is expressed in the spinal cord, prefrontal and frontal cortex, striatum, thalamus and amygdala; this pattern of expression is preserved throughout the development and adulthood [2, 3]. Its function is related to neuronal migration, dendritic arborisation and synaptic transmission [4]. The crucial role of *CNTNAP2* in the human brain became clear when Strauss *et al*, reported homozygous mutations in Old Order Amish families segregating with a severe Mendelian condition, described as cortical dysplasia-focal epilepsy (CDFE) syndrome (OMIM 610042) [5]. In 2009, additional patients with recessive mutations in *CNTNAP2* were reported, with clinical features resembling Pitt-Hopkins syndrome [6]. To date 33 patients, mostly from consanguineous families, have been reported with homozygous or compound deletions and truncating mutations in *CNTNAP2* [5-9], and are collectively described as having CASPR2 deficiency disorder [7]. The common clinical features in this phenotype include severe intellectual disability (ID), seizures with age of onset at two years and concomitant speech impairments or language regression. The phenotype is often accompanied by dysmorphic features, autistic traits, psychomotor delay and focal cortical dysplasia.

*CNTNAP2* is also thought to contribute to the phenotype in patients with interstitial or terminal deletions at 7q35 and 7q36. Interstitial or terminal deletions encompassing *CNTNAP2* and several other genes have been described in individuals with ID, seizures, craniofacial anomalies, including microcephaly, short stature and absence of language [10]. The severe language impairments observed in patients with homozygous mutations or karyotypic abnormalities involving *CNTNAP2* suggested a possible functional interaction with *FOXP2*, a gene for which heterozygous mutations lead to a monogenic form of language disorder [11]. Interestingly, *Vernes et al.,* found that the FOXP2 transcription factor has a binding site in intron 1 of *CNTNAP2*, regulating its expression [12]. Considering that a large proportion of autistic patients show language impairments and most individuals with homozygous mutations in *CNTNAP2* manifest autistic features, several studies investigated the potential involvement of *CNTNAP2* in autism spectrum disorder (ASD). In particular, two pioneering studies showed that single nucleotide polymorphism (SNP) markers rs2710102 and rs7794745 were associated with risk of ASD [13, 14]. Moreover, in subsequent studies, rs2710102 was implicated in early language acquisition in the general population [15], and showed functional effects on brain activation in neuroimaging studies [16-19]. Furthermore, genotypes at rs7794745 were associated with reduced grey matter volume in the left superior occipital gyrus in two independent studies [20, 21], and alleles of this SNP were reported to affect voice-specific brain function [22]. Genetic associations with ASD for these, and several other SNPs in *CNTNAP2*, have been reported in a number of studies [23-28]. Along with the first reports of variants associated with ASD, copy number variant (CNV) deletions have also been described in ID or ASD patients, which were proposed to be highly penetrant disease-causative mutations [13, 29-38]. To better understand the role of *CNTNAP2* in ASD pathophysiology, knockout mice were generated. Studies of these mice reported several neuronal defects when both copies of *CNTNAP2* are mutated: abnormal neuronal migration, reduction of GABAergic interneurons, deficiency in excitatory neurotransmission, and the delay of myelination in the neocortex [2, 39, 40].

These intriguing findings prompted additional investigations of *CNTNAP2* across other psychiatric disorders or language-related traits, with additional reports of variants being associated with schizophrenia (SCZ), bipolar disorder (BD), specific language impairment (SLI) and several other phenotypes or traits [12, 15, 41-50]. Consecutively, other studies reported CNV deletions in *CNTNAP2* in other psychiatric phenotypes such as schizophrenia [51, 52], bipolar disorder [52-54], and ADHD [55]; neurological disorders [56-61]; and language-related phenotypes [61-65]. Interestingly, several of these structural variants were found in intron 1 of *CNTNAP2*, encompassing the FOXP2 transcription factor binding site.

Our group recently performed CNV analysis in extended families with bipolar disorder [66], and found an intronic deletion in one individual which removed the FOXP2 binding site, prompting the need for a segregation analysis in this family. We therefore aimed in this current study to examine the evidence for a role of the *CNTNAP2* gene in multiple psychiatric phenotypes, performing a comprehensive analysis of common and rare variants, CNVs and *de novo* mutations using both in-house data and publically available datasets.

## RESULTS

### Examination of an intronic deletion in CNTNAP2 in an extended family with bipolar disorder

CNV microarray analysis was performed in two affected individuals from an extended family which included five relatives affected with bipolar I disorder. A drop in signal intensity for 340 consecutive probes was compatible with a deletion of 131 kb in intron 1 of *CNTNAP2* (hg19/chr7:146203548-146334635; Fig 1A). The deletion encompasses the described binding site for the transcription factor FOXP2 (hg19/chr7:146215016-146215040) [12]. The deletion was detected in one of the two affected individuals examined. To infer deletion segregation amongst relatives, WES-derived genotypes were used to create haplotypes across chromosome 7q35 (Fig 1B). The WES-derived haplotype analysis was uninformative due to incomplete genotype data (unaffected descendants of deceased patient 8404 not included in the WES study) and a likely recombination at 7q35 in the family. Thus experimental validation (in patient 8401) and CNV genotyping via quantitative PCR (qPCR) was performed in all individuals with DNA available from this family to assess the presence of the *CNTNAP2* intronic deletion. The deletion was validated in subject 8401, and was also detected in one unaffected descendant of deceased patient 8404 (Fig 1C), implying that this CNV would have been present in affected subject 8404, had DNA been available. The structural variant did not segregate with disease status in this family, and is unlikely to be a highly penetrant variant as it was observed also in an unaffected relative (Fig 1B).

**Fig 1.**
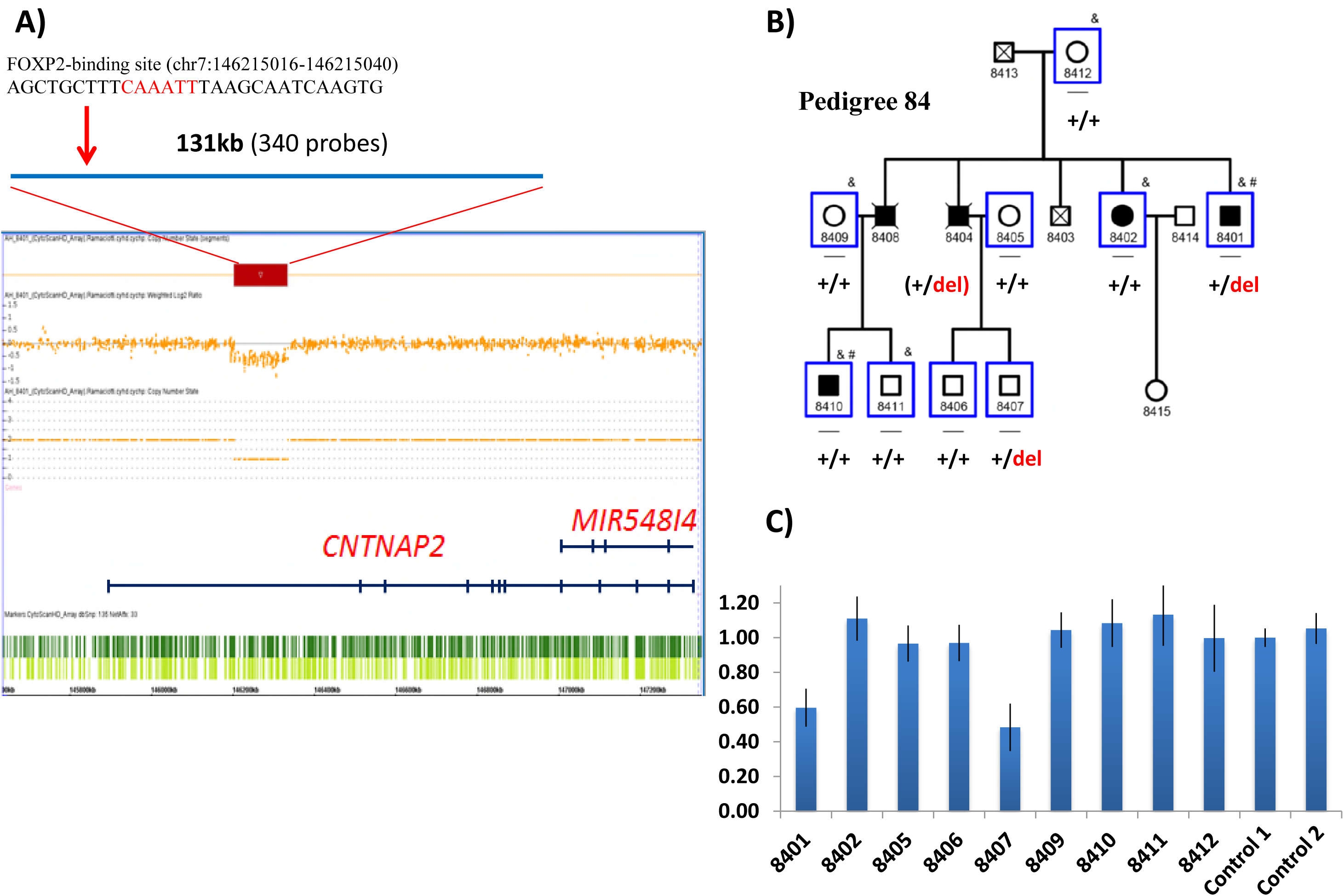
CNV deletion encompassing intron 1 of *CNTNAP2* in an extended family with bipolar disorder. A) CytoScan HD array output image shows the position of the drop in signal intensity of 340 probes, indicating a deletion spanning 131kb (chr7:146203548-146334635; GRCh37/hg19) found in the patient 8401. The position of the FOXP2 binding site within the deletion is shown above. B) The bipolar pedigree includes five patients with bipolar disorder I (BPI) across two generations. Symbols: _, individuals with DNA available; &, individuals with whole exome data; #, individuals analysed for genome-wide CNVs through the CytoScan HD array; blue squares, individuals included in CNV qPCR validation and genotyping analysis, for which heterozygous deletion carriers are indicated as “+/del” and non-carriers are indicated as “+/+”. Inferred genotypes are in parentheses. C) Gene dosage results of the qPCR experiments validating the deletion in patient 8401, and showing the deletion in unaffected subject 8407.

### Structural variants affecting CNTNAP2 amongst psychiatric phenotypes

Several deletions and duplications have been described in neuropsychiatric phenotypes thus far. In Fig 2, we present a comprehensive representation of all previously reported structural variants found in *CNTNAP2* in psychiatric disorders such as ASD or ID [13, 29-38], schizophrenia or bipolar disorder (including the 131kb deletion found in the present study) [51-54], ADHD [55], neurologic disorders such as epilepsy, Tourette syndrome or Charcot-Marie-Tooth [56-60]; and finally language-related phenotypes such as speech delay, childhood apraxia of speech and dyslexia [62-65]. Interestingly, the structural variants reported so far frequently map in intron 1, overlapping with the 131kb deletion found in our extended family, and extend in some cases up to exon 4. The distribution of those structural variants across different phenotypes does not vary with those found in control populations from the database of genomic variants (http://dgv.tcag.ca/dgv/app/home) (Fig 2), suggesting that structural variants in *CNTNAP2* are not rare events associated exclusively to disease but are present with rare frequency in the general population.

**Fig 2.**
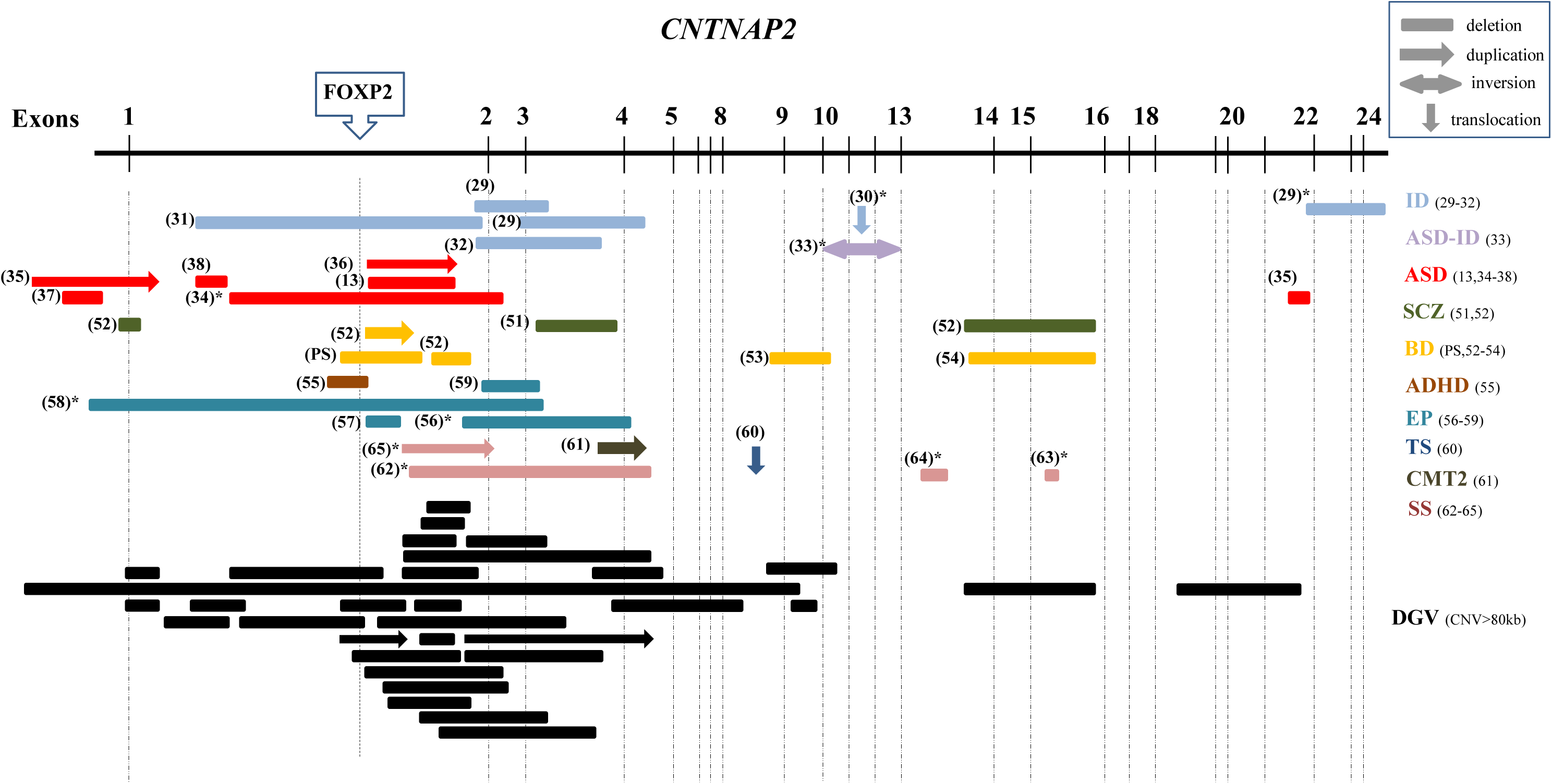
Overview of heterozygous CNVs spanning the *CNTNAP2* gene across several diseases. Abbreviations: ID (Intellectual disability), ASD (autism spectrum disorder), SCZ (schizophrenia), BD (bipolar disorder), ADHD (Attention-deficit/hyperactivity disorder), EP (epilepsy), TS (Tourette syndrome), CMT2 (axonal Charcot-Marie-Tooth), and SS (Speech spectrum: speech delay, childhood apraxia of speech and dyslexia). In parenthesis is reported the reference to each study. PS refers to this present study. ^∗^, additional rearrangements reported in this patient. The dashed lines represent the exons and the upper box shows the position of the FOXP2 binding site. In dark shading, CNVs≥80kb found in the general populations from the Database of Genomic Variants are shown.

### Analysis of CNTNAP2 common and rare variation in the susceptibility of psychiatric disorders

During the last decade, several association studies have been performed to assess the role of common variants of *CNTNAP2* in several psychiatric phenotypes. The functional relationship between *CNTNAP2* and the language-associated transcription factor FOXP2 prompted many studies to focus on language traits in autism or speech-related phenotypes [12-15, 23-28, 46-48, 50], but later, additional associations were also reported in other psychiatric phenotypes [41-45, 49]. In Table 1, we summarise all markers found significantly associated in these previous studies, and report the corresponding *P*-*value* from the Psychiatric Genomics Consortium GWAS for seven major psychiatric disorders: ADHD, anorexia nervosa, ASD, bipolar disorder, MDD, OCD and schizophrenia. Nominal associations were found with ASD for the following markers: rs802524 (*P*=0.016), rs802568 (*P*=0.008), rs17170073 (*P*=0.008), and rs2710102 (which is highly correlated with 4 SNPs: rs759178, rs1922892, rs2538991, rs2538976) (*P*=0.036); with schizophrenia for rs1859547 (*P*=0.044); with ADHD for rs1718101 (*P*=0.038); in MDD for rs12670868 (*P*=0.047), rs17236239 (*P*=0.006), rs4431523 (*P*=0.001); and with anorexia nervosa for rs700273 (*P*=0.013). The nominal association at rs1770073 and rs2710102 in ASD represents the only case in which the phenotype matches between the original report and the PGC dataset. The two SNPs rs7794745 and rs2710102, which were repeatedly reported as being associated and proposed to be functional SNPs, were not strongly associated with any phenotype (the most significant signal being *P*=0.036 for rs2710102 in autism). None of those associations survived corrections for multiple comparisons (Table 1).

**Table 1.**
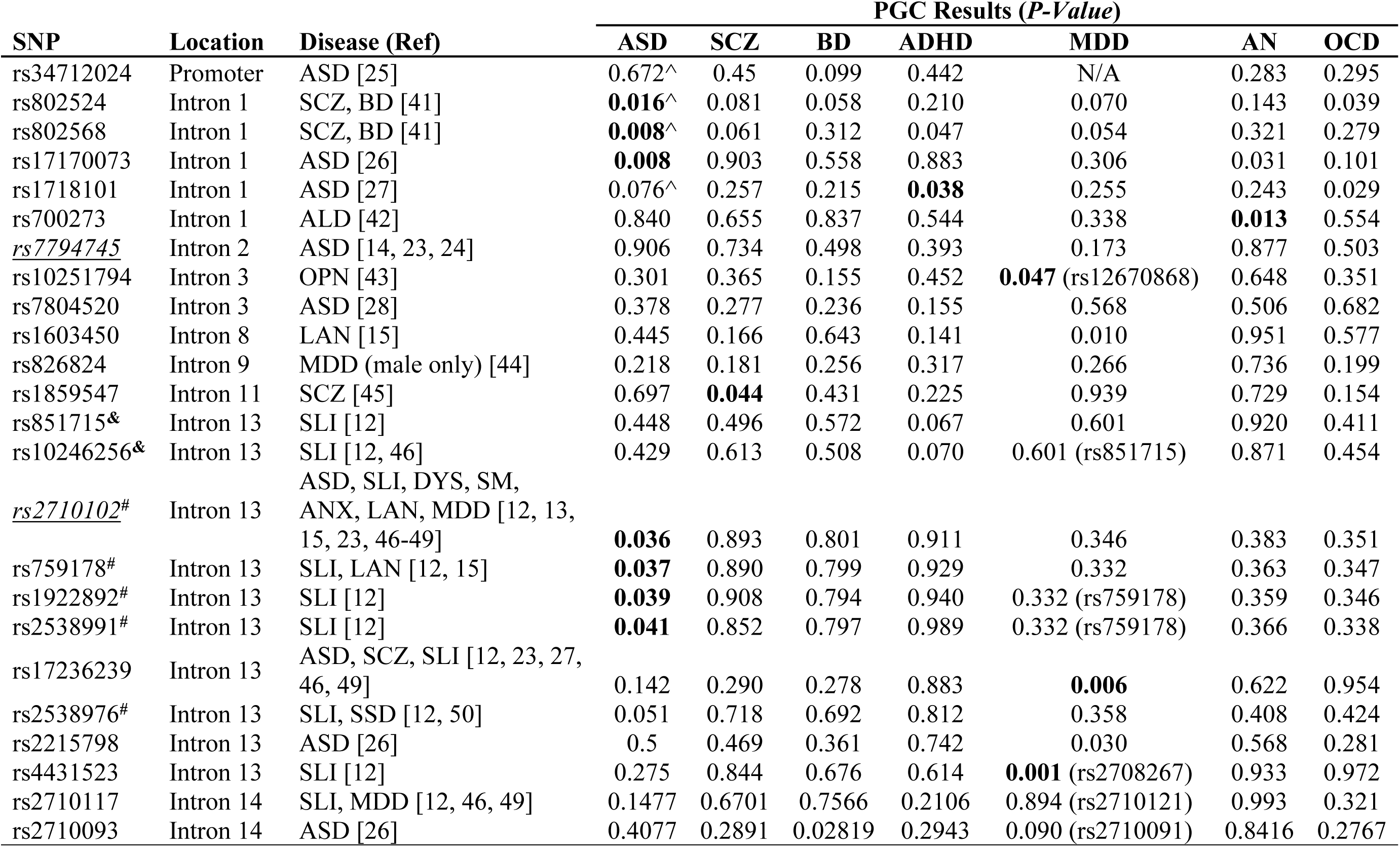
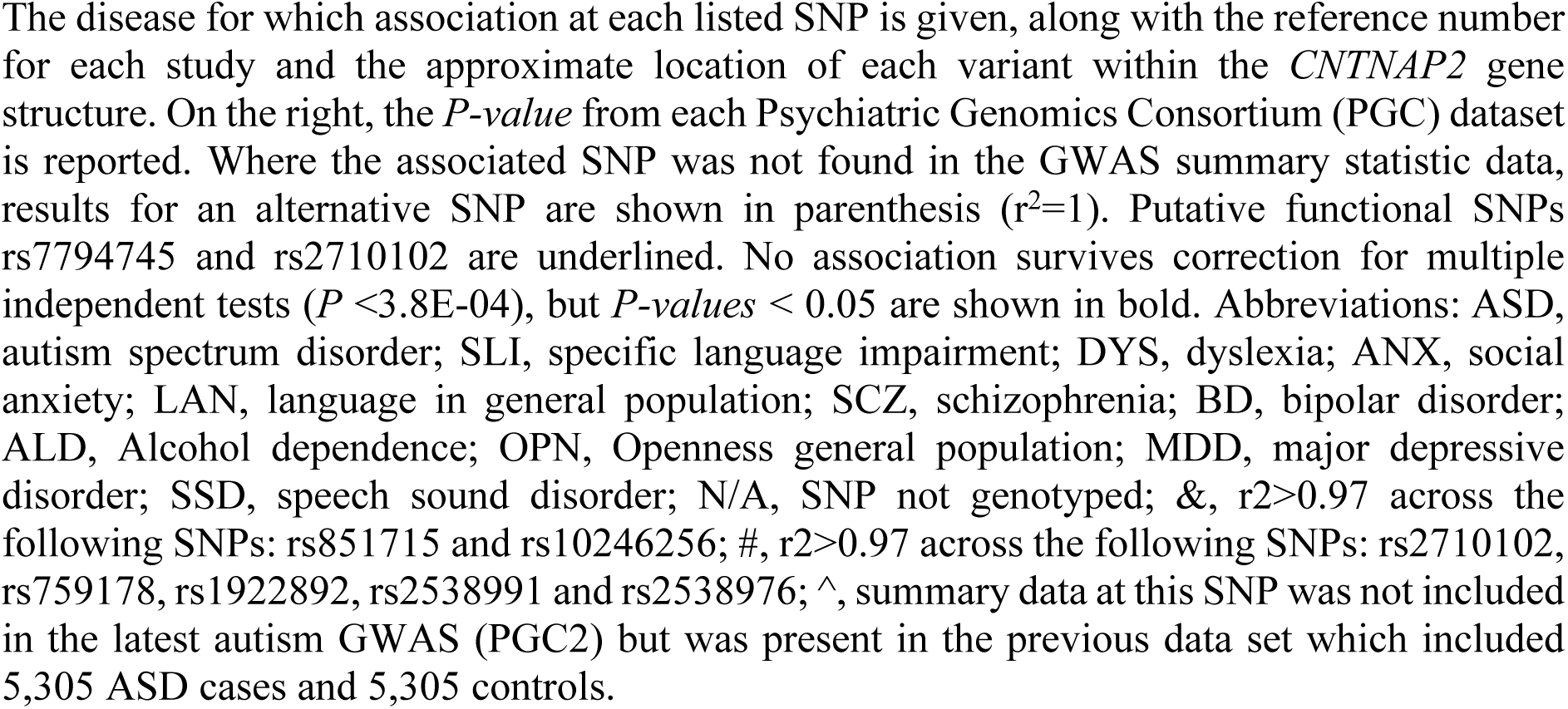
Common SNPs in *CNTNAP2* previously associated in psychiatric diseases, and their evidence for association in PGC datasets.

Next, we explored the contribution of common variants across *CNTNAP2* by performing a gene-based association study in European populations using GWAS summary statistics from PGC data of seven psychiatric disorders (Table 2).

**Table 2.** Gene-based tests for association of *CNTNAP2* across seven psychiatric disorders using GWAS summary statistics of the PGC data sets.

| Disease | N Cases – Controls | N SNPs tested | Gene-based *P*-value | Top SNP | Top SNP *P*-value |
| --- | --- | --- | --- | --- | --- |
| ADHD<sup>a</sup> | 19,099 - 34,194 | 7,538 | 0.16 | rs370840971 | 4.8E-04 |
| AN<sup>a</sup> | 3,495 - 10,982 | 9,318 | 0.33 | rs138287908 | 8.5E-05 |
| ASD<sup>a</sup> | 6,197 - 7,377 | 5,946 | 0.54 | rs1089600 | 0.0018 |
| BD<sup>a</sup> | 20,352 - 31,358 | 11,345 | 0.34 | rs181471483 | 3.6E-04 |
| MDD<sup>b</sup> | 9,240 - 9,519 | 1,214 | **0.029** | rs4725752 | 9.3E-04 |
| OCD<sup>a</sup> | 2,688 - 7,037 | 8,631 | 0.30 | rs6976859 | 8.7E-05 |
| SCZ<sup>a</sup> | 33,640 - 43,456 | 12,264 | 0.11 | rs78093069 | 1.1E-04 |

The numbers (N) of cases and controls in each dataset examined are given, along with the number of SNPs tested in each dataset. The name and *P*-value of the SNP with most significant association is given. Abbreviations: ADHD, attention-deficit/hyperactivity disorder; AN, anorexia nervosa; ASD, Autism spectrum disorder; BD, bipolar disorder; MDD, major depressive disorder; OCD, obsessive compulsive disorder; SCZ, schizophrenia; <sup>a</sup>, European individuals from the PGC2 data sets; <sup>b</sup>, European individuals from the PGC1 data sets.

The test included a dense coverage of SNPs across *CNTNAP2:* from 1,214 SNPs in MDD up to 12,264 SNPs in schizophrenia. The results suggest that common variants overall do not contribute to disease susceptibility of these phenotypes (Table 2). The most significant association observed was for MDD phase 1 analysis (*P*=0.029), which is the dataset with the most modest coverage of markers.

In our final analysis of the PGC datasets, we selected 63 predicted functional SNPs in *CNTNAP2* and performed a cross-disorder meta-analysis, aimed to test evidence for association with common functional variants across psychiatric disorders. Nominal significance of association was observed for 11 predicted functional SNPs with *P*-*values* ranging from 0.01 and 0.05, but none survive correction for multiple comparisons (Table 3).

**Table 3.**
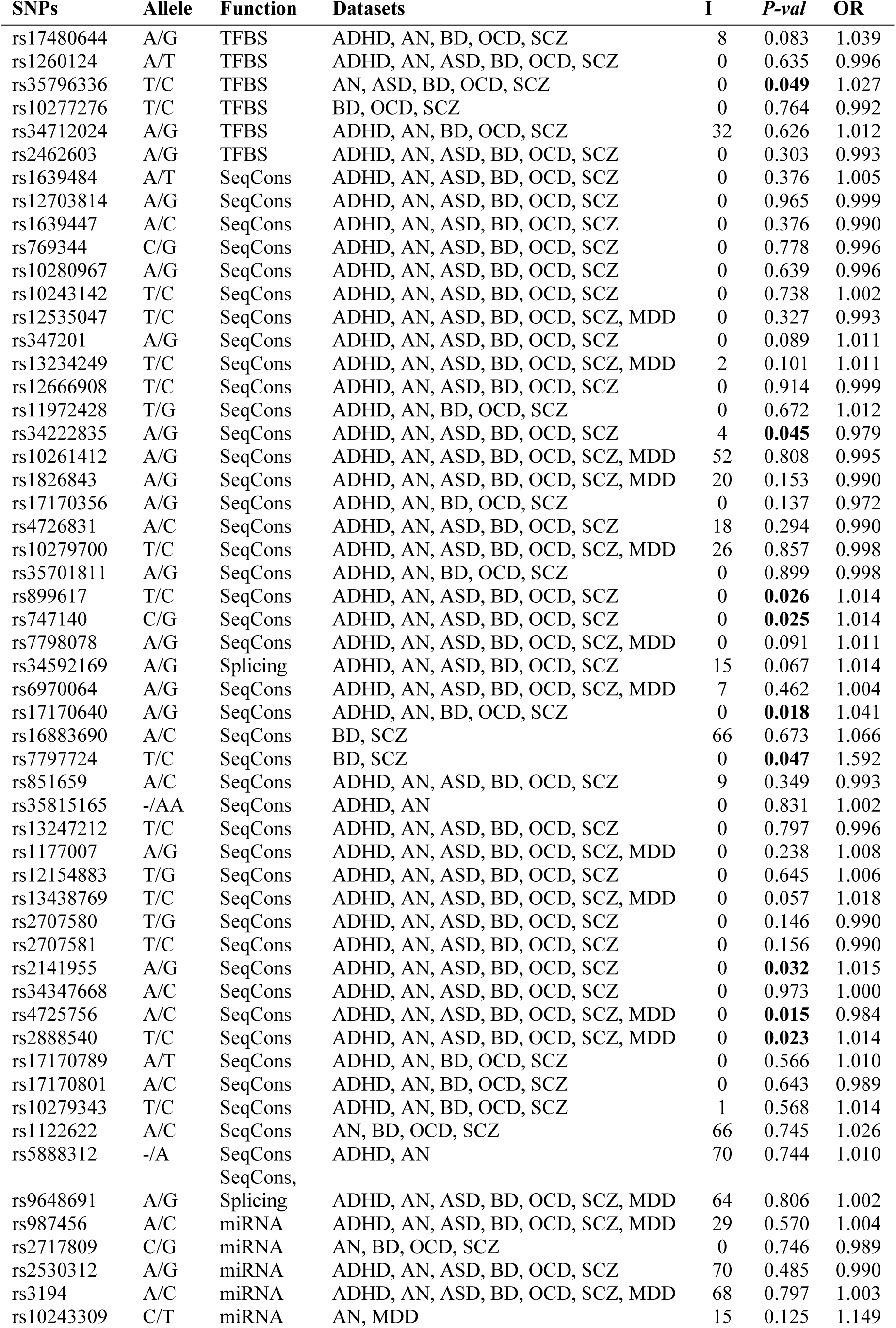
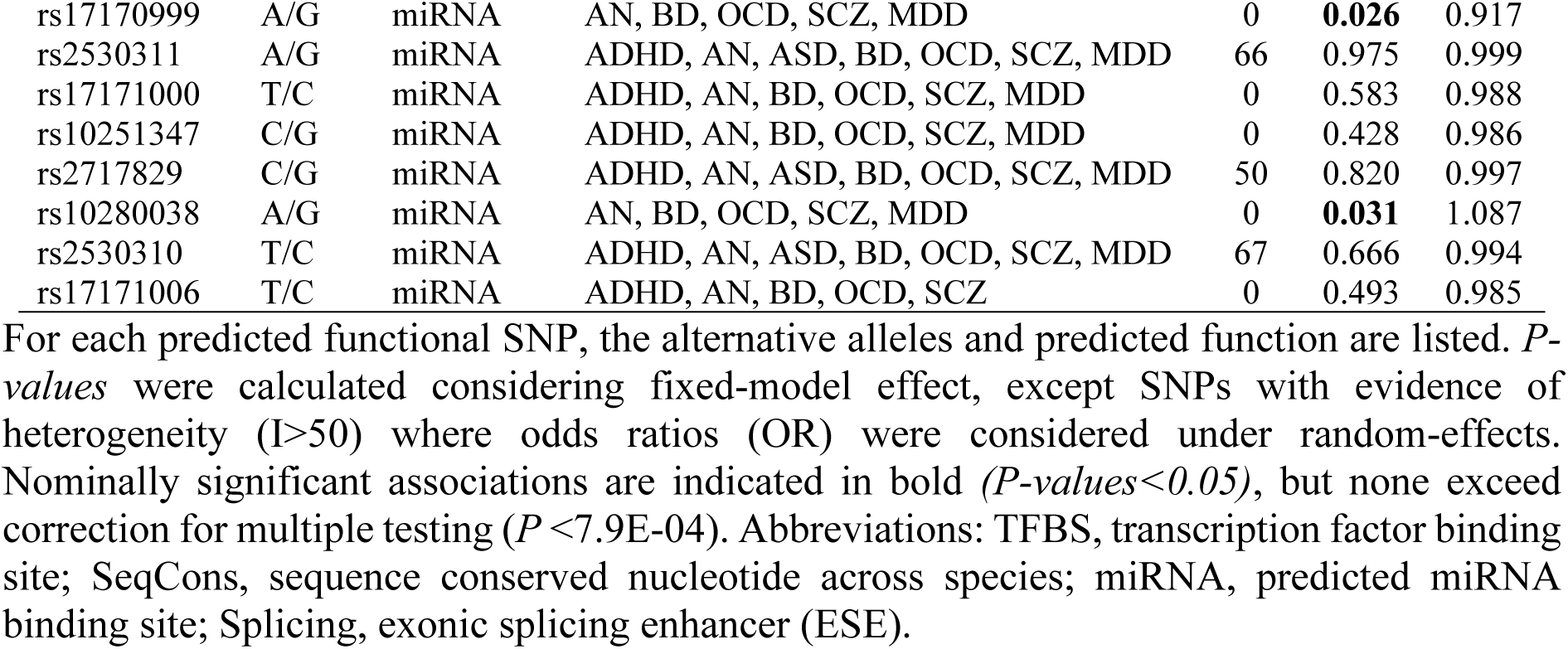
Cross psychiatric disorders meta-analysis of predicted functional SNPs.

The only SNP predicted to be functional and which was previously reported as being associated with autism was rs34712024 (Table 2) [25], but this variant was not associated with autism in PGC dataset (*P*=0.67), nor other psychiatric phenotypes examined (Table 2).

*De novo* variants in protein-coding genes which are predicted to be functionally damaging are considered to be highly pathogenic and have been extensively explored to implicate genes in psychiatric diseases, especially in ASD and schizophrenia [67]. We explored publically available sequence data from previous projects in psychiatric disorders to assess the rate of coding *de novo* variants in *CNTNAP2* using two databases (NPdenovo, http://www.wzgenomics.cn/NPdenovo/; and denovo-db, http://denovo-db.gs.washington.edu/denovo-db/). No truncating or missense variants were identified across *CNTNAP2* in 15,539 families (including 2,163 controls), and synonymous variants were reported in only two probands with developmental disorder (Table 4).

**Table 4.** *CNTNAP2 de novo* variants identified across several disease-specific sequencing projects.

| Phenotype | N Families | Intronic | Synonymous | Missense |
| --- | --- | --- | --- | --- |
| ASD | 6,171 | 106 | - | - |
| SCZ | 1,164 | - | - | - |
| EE | 647 | - | - | - |
| ID | 1,101 | - | - | - |
| DD | 4,293 | - | 2 | - |
| Controls | 2,163 | 13 | - | - |

The number (N) of families in each dataset examined is given. The full list of *de novo* variants observed is listed in S2 Table. Abbreviations: ASD, autism spectrum disorder; SCZ, schizophrenia; EE, epilepsy; ID, intellectual disability; DD, developmental disability.

Finally, we explored the potential impact of pathogenic ultra-rare variants (URV) in *CNTNAP2* using available sequencing datasets of 4,483 patients with ASD and 6,135 patients with schizophrenia compared with 13,042 controls. We considered only those variants predicted to be pathogenic in both SIFT and Polyphen and which are ultra-rare (MAF<0.0001 in Non-Finnish European population). No difference in the total number of URV was observed between ASD and controls (*P*=0.11), or between schizophrenia patients and controls (*P*=0.78) (Table 5).

**Table 5.** Burden analysis of *CNTNAP2* ultra-rare variants (URVs) in ASD and SCZ.

|  | N Individuals | N Pathogenic URVs | *P-Value* |
| --- | --- | --- | --- |
| Controls | 13,042 | 59 |  |
| SCZ | 6,135 | 26 | 0.78 |
| ASD | 4,483 | 29 | 0.11 |

The selection of variants included missense variants which are predicted to be pathogenic, truncating variants and canonical splice-site variants. The full list of URVs observed is provided in S3 Table. Abbreviations: SCZ, schizophrenia; ASD, autism spectrum disorder

## DISCUSSION

During the last decade, the *CNTNAP2* gene has received considerable attention in the psychiatric genetics field, with a large body of studies examining gene dosage, common or rare variants across multiple major psychiatric disorders, which together provided compelling evidence that *CNTNAP2* may be a risk gene with pleiotropic effects in psychiatry. While homozygous mutations in this gene lead to a rare and severe condition described as CASPR2 deficiency disorder (CDD) [7], characterized by profound intellectual disability, epilepsy, language impairment or regression [7, 8], heterozygous mutations or common variants have been suggested to be implicated in autism, whose features overlap with some observed in CDD. *CNTNAP2* is categorised in the SFARI database (https://gene.sfari.org) as a strong candidate gene for ASD (category 2.1). Heterozygous deletions encompassing the *CNTNAP2* gene were described not only in autism but also in a wide range of phenotypes, including psychiatric or neurologic disorders, and language-related deficiencies. These structural variants were generally described as causative or highly penetrant [13, 29, 31, 55, 57, 59].

Here, we describe a new deletion in a bipolar disorder patient encompassing intron 1 of *CNTNAP2*, which overlaps with structural variants described in a number of other psychiatric patients. This heterozygous deletion, which removes the FOXP2 transcription factor binding site, was found in an individual with bipolar I disorder from an extended family with five affected members. This deletion was observed in only two of the affected relatives and was absent from two affected relatives, but was also observed in one unaffected relative who underwent diagnostic interview at age >40 and therefore beyond the typical age of symptom onset. Hence, the deletion was not segregating with the disease and is unlikely to represent a highly penetrant risk variant in this family. Examination of the distribution of all structural variants described thus far in psychiatric or neurologic patients showed comparable mapping with those found in the general population, suggesting that structural variants affecting *CNTNAP2* may be less relevant in disease susceptibility than previously considered. Eleven CNVs are described in the general population against sixteen expected (z=0.43) in ExAC database (http://exac.broadinstitute.org), and the haploinsufficiency score (0.59) is relatively moderate [68], suggesting that *CNTNAP2* has a moderate tendency to be intolerant to structural variants. However, a case-control CNV analysis is needed in psychiatric disorders, but would require a very large sample due to the rarity of CNVs at this locus. A close clinical psychiatric examination of the 66 parents with heterozygous deletions across *CNTNAP2* of CDD provide information on the prevalence of psychiatric conditions in individuals carrying *CNTNAP2* CNVs. All heterozygous family members carrying deletions or truncating mutations were described as phenotypically healthy, suggesting a lack of correlation between these deletions and any major psychiatric condition. Furthermore, parents who were carriers for heterozygous deletions in psychiatric/neurologic patients were described as unaffected at the time of reporting [13, 29, 31, 37, 54, 62], with the exception of one father of a proband with neonatal convulsion or another father of an epileptic patient reported as affected [56, 59]. Moreover, discordant segregation for deletions in *CNTNAP2* was also observed in an ASD sib-pair [13]. Several psychiatric patients who were reported to carry heterozygous structural variants in *CNTNAP2* were also described with translocations or other chromosomal abnormalities [29, 30, 33, 34, 56, 58, 62-65], therefore it is possible that these aberrations may explain the phenotype independently from the observed CNVs in *CNTNAP2*.

*CNTNAP2*^-/-^ knock-out mice have been proposed as valid animal model for ASD considering the phenotypic similarities between ASD and the CASPR2 deficiency disorder [2]. *CNTNAP2*^-/-^ knock-out mice showed abnormalities in the arborisation of dendrites, maturation of dendritic spines, defects in migration of cortical projection neurons, and reduction of GABAergic interneurons [2, 4]. Controversially, ASD is not a core feature in the most recent patient series reported with CASPR2 deficiency disorder [7, 8]. The association previously proposed around the relationship between heterozygous deletions in *CNTNAP2* and ASD does not have a support from mouse models, as heterozygous mice did not show any behavioural or neuropathological abnormalities that were observed in homozygous knockouts [2]. Notwithstanding this, it is possible that the combination of heterozygous *CNTNAP2* deletions in a genomic background of increased risk (through inheritance of other common and rare risk variants at other loci) may lead to psychiatric, behavioural or neuropathological abnormalities.

Common variants in *CNTNAP2* are another class of genetic variation associated with several psychiatric or language-related phenotypes. The most interesting finding from these studies converge on markers rs7794745 and rs2710102, originally reported in ASD [13, 14], and replicated later in ASD or implicated in other phenotypes [12, 15, 23, 24, 46-48]. Neuroimaging studies have supported the notion that these common variants play a role in psychiatric disorders. SNP rs2710102 has been implicated in brain connectivity in healthy individuals [16, 18, 19], and rs7794745 was implicated in audio-visual speech perception [69], voice-specific brain function [22], and was associated with reduced grey matter volume in left superior occipital gyrus [20, 21]. These studies focused principally on language tasks in general population, given the reported suggestive implications of *CNTNAP2* in language impairment traits of ASD or language-related phenotypes. However, the direct role of *CNTNAP2* in language is still unclear; indeed the language regression observed in patients with CASPR2 deficiency are concomitant with seizure onset and may represent a secondary phenotypic effect caused by seizures [7]. On the other hand, the first genetic association of rs7794745 and rs2710102 with ASD, as well as the other psychiatric diseases were based in studies with limited sample size, and recent studies failed to replicate associations between the two markers and ASD [70, 71]. Individual alleles associated in the past with limited numbers of patients warrant replications in adequately powered samples to ascertain *bona fide* findings considering the small size effects of common variants [72], which we attempted here using the largest case-control cohorts currently publicly available (PGC datasets). We did not find evidence for significant association of previous reported common variants, nor did we find functional SNPs with a role across disorders, or observe a combined effect for common variants of *CNTNAP2* in the susceptibility of psychiatric disorders.

Rare variants of *CNTNAP2* both in the promoter or coding region were also reported to play a role in the pathophysiology of ASD [25, 33]. A recent study including a large number of cases and controls did not find association of rare variants of *CNTNAP2* in ASD [73]. Here we report the largest sample investigated thus far in ASD and schizophrenia for rare variants in *CNTNAP2*, which suggest that rare variants from this gene do not play a major role in these two psychiatric disorders. Furthermore, the identification of *de novo* variants in *CNTNAP2* in combined psychiatric sequencing projects of over 15,500 trios suggest that *de novo* variants in this gene do not increase risk for psychiatric disorders.

While functional studies show a relationship between certain deletions or rare variants of *CNTNAP2* with neuronal phenotypes relevant to psychiatric illness [25, 54, 74], we show that the genetic link between these variants and psychiatric phenotypes is tenuous. However, this does not dispel the evidence that the *CNTNAP2* gene, or specific genetic variations within this gene, may have a real impact on neuronal functions or brain connectivity.

Nowadays we are able to combine large datasets to ascertain the real impact of candidate genes described in the past in psychiatric disorders. Here we performed analyses using large publically available datasets investigating a range of mutational mechanisms which impact variability of *CNTNAP2* across several psychiatric disorders. In conclusion, our results converge to show a limited or likely neutral role of *CNTNAP2* in the susceptibility of psychiatric disorders.

## MATERIALS AND METHODS

### Extended family with bipolar disorder and CNV in CNTNAP2

The extended family presented here (Fig 1B) provides a molecular follow-up from a previously reported whole exome sequencing (WES) study of multiplex BD families, augmented with CNV microarray data [66]. This multigenerational pedigree, was collected through the Mood Disorders Unit and Black Dog Institute at the Prince of Wales Hospital, Sydney, and the School of Psychiatry (University of New South Wales in Sydney) [75-79]. Consenting family members were assessed using the Family Interview for Genetic Studies (FIGS) [80], and the Diagnostic Interview for Genetic Studies (DIGS) [81]. The study was approved by the Human Research Ethics Committee of the University of New South Wales, and written informed consent was obtained from all participating individuals. Blood samples were collected for DNA extraction by standard laboratory methods. Three of the five relatives with bipolar disorder type I (BD-I) had DNA and WES-derived genotype data available, and six unaffected relatives with DNA and WES data were available for haplotype phasing and segregation analysis (Fig 1B).

Genome-wide CNV analysis was performed via CytoScan^®^ HD Array (Affymetrix, Santa Clara, CA, USA) in 2 distal affected relatives (individuals 8410 and 8401; Fig 1B), using the Affymetrix Chromosome Analysis Suite (ChAS) software (ThermoFisher, Waltham, MA, USA). Detailed information on CNV detection and filtering criteria have been previously described [66]. We identified a 131kb deletion in intron 1 of *CNTNAP2* in individual 8401. WES-derived genotypes were used for haplotype assessment to infer CNV segregation amongst relatives, as previously described [66]. Next, we experimentally validated the *CNTNAP2* CNV via quantitative PCR (qPCR) in all available family members. Validation was performed in quadruplicate via a SYBR Green-based quantitative PCR (qPCR) method using two independent amplicon probes, each compared with two different reference amplicon probes in the *FOXP2* and *RNF20* genes (S1 Table). Experimental details are available upon request.

### Common variant association in CNTNAP2 using publically available datasets

We sought to replicate previously reported *CNTNAP2* SNP associations in a range of psychiatric phenotypes or traits using GWAS summary-statistic data of the Psychiatric Genomics Consortium (https://med.unc.edu/pgc/results-and-downloads).

Firstly, we report the corresponding *P*-*values* of specific previously associated markers for case-control cohorts with autism spectrum disorder (ASD), schizophrenia (SCZ), bipolar disorder (BD), attention-deficit hyperactivity-disorder (ADHD), major depressive disorder (MDD), anorexia nervosa (AN), and obsessive compulsive disorder (OCD). If a specific SNP marker was not reported in an individual GWAS dataset, we selected another marker in high linkage disequilibrium (r^2^~1, using genotype data from the CEU, TSI, GBR and IBS European populations in 1000genomes project; http://www.internationalgenome.org).

Next, a gene-based association for common variants was calculated with MAGMA [82], using variants within a 5 kb window upstream and downstream of *CNTNAP2*. Selected datasets were of European descent, derived from GWAS summary statistics of the Psychiatric Genomics Consortium (https://med.unc.edu/pgc/results-and-downloads): SCZ (33,640 cases and 43,456 controls), BD (20,352 cases and 31,358 controls), ASD (6,197 and 7,377 controls), ADHD (19,099 cases and 34,194 controls) and MDD (9,240 cases and 9,519 controls) [83-87]. Analyses were performed combining two different models for higher statistical power and sensitivity when the genetic architecture is unknown: the combined *P*-*value* model, which is more sensitive when only a small proportion of key SNPs in a gene show association; and the mean SNP association, which is more sensitive when allelic heterogeneity is greater, and a larger number of SNPs show nominal association.

Finally, we selected SNPs predicted to be functional within a 5kb window upstream/downstream of *CNTNAP2* (e.g. located in transcription factor binding sites, miRNA binding sites etc; https://snpinfo.niehs.nih.gov), and assessed a potential cross-disorder effect using GWAS summary statistics data of the PGC by performing a meta-analysis in PLINK [88]. The Cochran’s Q-statistic and I^2^ statistic were calculated to examine heterogeneity among studies. The null hypothesis was that all studies were measuring the same true effect, which would be rejected if heterogeneity exists across studies. For all functional SNPs, when heterogeneity between studies was I>50% (*P*<0.05), the pooled OR was estimated using a random-effects model.

### Analysis of rare variants in CNTNAP2 in ASD and schizophrenia, and de novo variants across psychiatric cohorts

The impact of rare variants of *CNTNAP2* was assessed using sequencing-level data from the following datasets: WES from the Sweden-Schizophrenia population-based Case-Control cohort (6,135 cases and 6,245 controls; dbGAP accession: phs000473.v2.p2); ARRA Autism Sequencing Collaboration (490 BCM cases, BCM 486 controls, and 1,288 unrelated ASD probands from consent code c1; dbGAP accession: phs000298.v3.p2); Medical Genome Reference Bank (2,845 healthy Australian adults; https://sgc.garvan.org.au/initiatives/mgrb); individuals from a Caucasian Spanish population (719 controls [89, 90]); in-house ASD patients (30 cases; [91]); and previous published data set in ASD (2,704 cases and 2,747 controls [73]). The selection of potentially etiologic variants was performed based on their predicted pathogenicity (missense damaging in both SIFT and polyphen 2, canonical splice variants, stop mutation and indels) and minor allele frequency (MAF<0.0001 in non-Finnish European populations using the Genome Aggregation Database; http://gnomad.broadinstitute.org/). A chi square statistic was used to compare separately the sample of schizophrenia patients (6,135 cases) and the combined ASD data sets (4,512 cases) with the combined control data sets (13,042 individuals).

Two databases for *de novo* variants were used to identify *de novo* variants in *CNTNAP2* [92, 93], which comprise data for the following samples: autism spectrum disorder (6,171 families), schizophrenia (1,164 families), epilepsy (647 families), intellectual disability (1,101 families), developmental disorders (4,293 families) and controls (2,163).

## ACKNOWLEDGMENTS

This study was funded by the Australian National Medical and Health Research Council (NHMRC) Project Grant 1063960 and supported by NHMRC Project Grant 1066177, and Program Grant 1037196. We gratefully acknowledge the Janette Mary O’Neil Research Fellowship (to JMF) and Mrs. Betty Lynch for supporting this work. DNA for the bipolar disorder family was extracted by Genetic Repositories Australia (GRA; www.neura.edu.au/GRA), an Enabling Facility that was supported by NHMRC Enabling Grant 401184. DNA samples were prepared and whole exome sequenced at the Lottery State Biomedical Genomics Facility, University of Western Australia. We would also thank Xose S. Puente from the University of Oviedo (Spain) for providing us the data from a Spanish control population. We are grateful to all participants and their families, as well as clinical collaborators who were originally involved in collecting and phenotyping these families, including Laila Tabassum, and Adam Wright (UNSW).

## SUPPORTING INFORMATION

**S1 Table. Primers used in the CNV validation for the *CNTNAP2* intronic deletion.**

**S2 Table. Full list of *de novo* variants in *CNTNAP2* gene.**

**S3 Table. Full list of Ultra-Rare Variants (URVs) in available sequencing datasets**.

